# *Canavalia ensiformis* intercrop to reduce weeds and stalk borer damage in sugarcane

**DOI:** 10.64898/2026.01.26.701649

**Authors:** Mansuy Alizé, Christina Mathias, Martin José, Marion Daniel, Chabalier Maxime, Nibouche Samuel

## Abstract

The aim of this study was to assess the efficiency of two intercrop species, *Canavalia ensiformis* and *Desmodium intortum*, to reduce weed growth, herbicide use and damage by the stalk borer *Chilo sacchariphagus* in sugarcane cropping system in Reunion.

We compared six inter-row management techniques: four treatments combined the two intercrops *Canavalia ensiformis* or *Desmodium intortum* sown either early (between 0 and 2.1 months after sugarcane emergence) or late (between 1.3 and 3.7 months after sugarcane emergence), a treatment where no weeding was carried out on the inter-row, and a treatment with chemical weeding of the inter-row (CcWp). In all treatments, the sugarcane row was weeded chemically and manually. The six treatments were compared in a multilocal randomized block design with three localities, during one or two crop cycles depending on the locality.

*Desmodium intortum* produced poor ground coverage in half of the trial x crop cycles and was therefore found unsuitable for use as an intercrop of sugarcane in our conditions. On the opposite, *Canavalia ensiformis* quickly covered the inter-rows, regardless of the sowing date. The *Canavalia ensiformis* intercrops allowed a reduction of herbicide consumption by 63% when sown early and by 28% when sown late, compared to the CcWp control treatment. Both *Canavalia ensiformis* intercrops caused a reduction of weed coverage on the inter-row similar or better than the chemical control. However, the early sown *Canavalia ensiformis* intercrop caused a 18.6 t.ha^-1^ yield loss compared the chemical control. No yield loss was detected with the late sown intercrop. A significant reduction of stalk damage by a 0.8-fold factor was observed in the early sown *Canavalia ensiformis* treatment.

## 1. Introduction

Sugarcane represents a major source of sugar, ethanol and material for electric energy productions in tropics and sub-tropics. In the energy sector, the demand for ethanol and electricity is likely to increase, as it could replace fossil fuels and reduce greenhouse gas emissions (Goldemberg et al., 2014; Leal et al., 2013). Grown mainly in the tropics, sugarcane plantations faced various harmful organism such as insects (white grubs and stem borers), fungal or bacterial diseases (smut, leaf scald, gumming) as well as weeds (Marnotte et al. 2010). Therefore, similarly to other crops, large amounts of chemical inputs are used to prevent yield losses (Oerke and Dehne, 2004).

In Reunion Island, sugarcane plays an important economic, societal and environmental role covering 54% of agricultural land (Agreste, 2023). Thanks to the use of varietal resistance to diseases and biological control of white grubs, no synthetic insecticides or fungicides are registered for use in sugarcane crops on the island. Nevertheless, two major threats remain, weeds and stem borers. Despite being considered a giant grass, sugarcane has a relatively slow growth rate during the first three months, which make it sensitive to competition with weeds (Marion and Marnotte, 1991). Consequently, herbicides are still recommanded to control weeds (Antoir et al., 2016). On its side, the stem borer *Chilo sacchariphagus* is considered causing losses up to 25% of the yield in some situations (Goebel et al. 1999), although it is subject to partial varietal control (Nibouche and Tibère 2009, 2010). To meet new societal demands, there is an urgent need to reduce the use of chemical pesticides.

In tropical agrosystems, the use of cover or companion plants to control weeds can be one of the solutions to help reduce herbicide inputs (Bhaskar et al., 2018; Mennan et al., 2020; Ranaivoson et al., 2018) before planting (Lu et al., 2000) or during crop growth as intercrop (Vandermeer, 1992). These plants are being increasingly used in innovative cropping systems to favor biological regulation and to deliver agro-ecosystem services such as weed or pest control (Altieri et al., 2011; Snapp et al., 2005). Consequently, the use of cover plants in rotation or intercropping with sugarcane is an increasing practice worldwide (Soares et al., 2017 in Brazil; Ali et al., 2018 in Africa; Paungfoo-Lonhienne et al., 2017 in Australia; Hemwong et al., 2009 in Asia) but is still recently investigated in Reunion (Christina et al., 2018; Mansuy et al., 2019).

To answer both weed and stem borer threats in Reunion, cover plants could be used with a push-pull approach (Cook et al. 2007). The strategy is based on the combined use of a repulsive plant present in the plot and grown in association with the crop and an attractive plant located at the edge of the field. Such strategy has been sucessfully used in Kenya to control the maize stem borer *Chilo partellus*, with *Desmodium* species (Khan et al. 2001). In Reunion, *D. intortum* is used as a cover plant in coffee plantations and is investigated in intercropping with sugarcane to control weeds. Moreover, another promising cover plant species to control weeds in Reunion is the jack bean *Canavalia ensiformis* (Mansuy et al., 2019). Some references indicate that *C. ensiformis* could reduce crop damage from certain insects, including the African sugarcane borer *Eldana saccharina* (Agboka, 2009) and the coffee weevil *Hypothenemus hampei* (Pohlan, et al 2008).

The aim of this study was to assess the efficiency of two intercrop species, *Canavalia ensiformis* and *Desmodium intortum*, to reduce i) weed growth, ii) herbicide use and iii) stalk borer damage in sugarcane cropping system in Reunion.

## 2. Material and Methods

### 2.1. Trial implantation

Three field trials were carried out between 2013 and 2016 in Reunion Island (South-West Indian Ocean) in contrasted localities. One trial was carried out in an experimental research station located in the north of the island and two trials were carried out in farmer fields in the East and North-West (Table 1). The Saint Paul trial was monitored during one crop cycle while the two other trials were monitored during two crop cycles.

**Table 1.** Localisation and characteristics of the trials. Soil classification according to IUSS Working Group WRB (2015).

|  | Locality |  |  |
| --- | --- | --- | --- |
|  | Sainte Marie | Saint Paul | Saint Benoît |
| Location | North<br>Latitude : -20.90<br>Longitude : 55.53 | North-West<br>Latitude : -21.07;<br>Longitude : 55.28 | East<br>Latitude : -21.05<br>Longitude : 55.69 |
| Elevation | 68 m | 545 m | 150 m |
| Monitored crop cycles | 2014-2015 (planted cane)<br>and 2015-2016 (first ratoon) | 2014-2015 (first ratoon) | 2013-2014 (second ratoon)<br>and 2014-2015 (third ratoon) |
| Soil | Nitisol | Andic umbrisol | Nitisol |
| Irrigation | sprinkler | sprinkler | none |
| 10-years mean annual pluviometry | 1 449 mm | 886 mm | 2953 mm |
| 10-years average temperature | 23.95 °C | 19.82 °C | 22.69 °C |

### 2.2. Treatments

Six treatments were compared in all trials and included four different cover crop, one control plot with full chemical weed control and one treatment with weed control on the row only (Table 2). Each trial was arranged as a randomized complete block design with three replications (Saint Paul) or four replications (Sainte Marie and Saint Benoît). Each elementary plot consisted of four rows, 10 m-long and 1.5 m spaced. Fertilization (Supplementary Table S1) and irrigation were similar to recommended practices for the area of implantation of each trial.

**Table 2.**
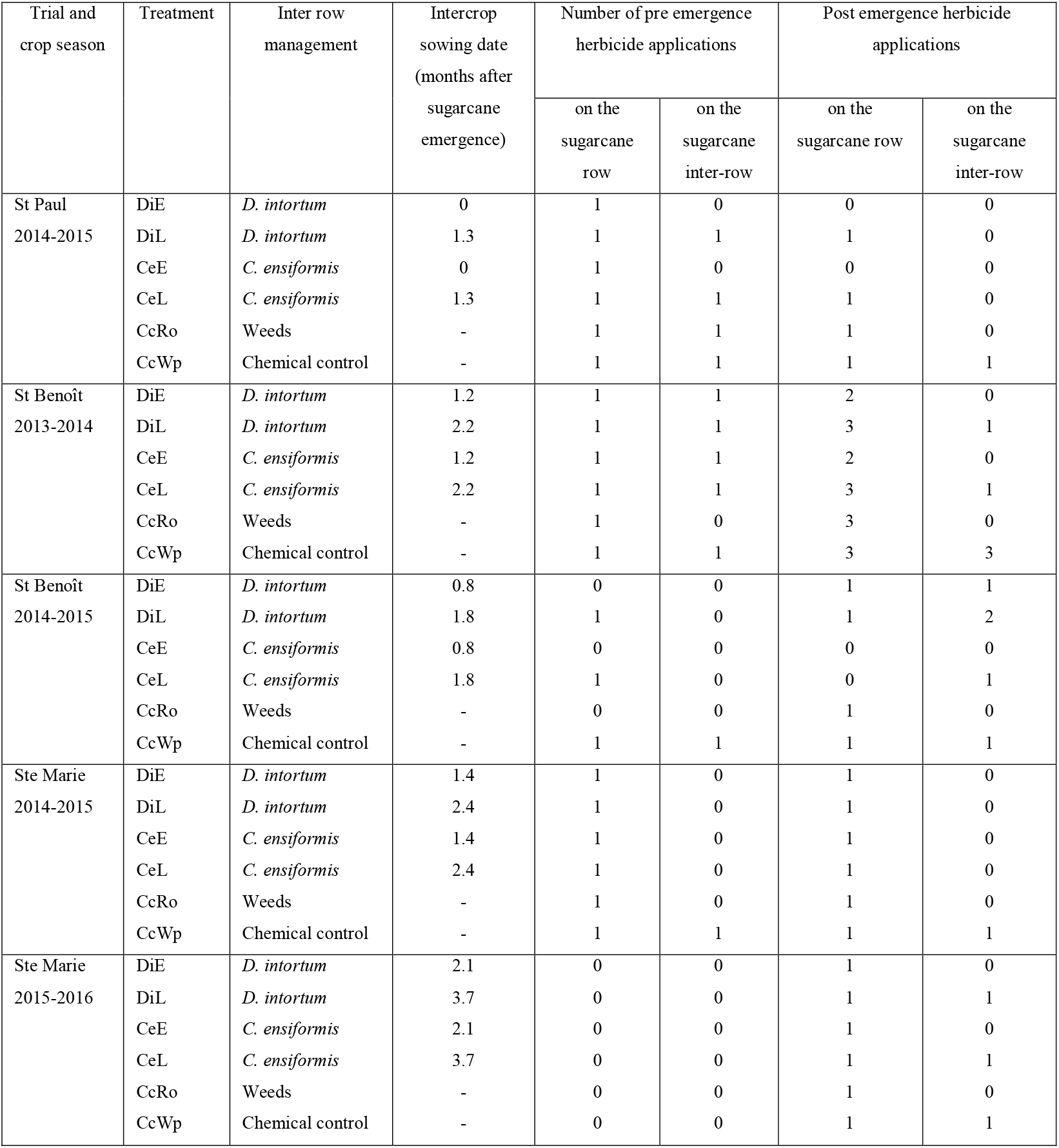
Compared treatments: type of inter-row management, date of sowing of intercrops, number of herbicide applications. DiE = D. intortum sown early; DiL = D. intortum sown late; CeE = C. ensiformis sown early; CeL = C. ensiformis sown late; CcRo = chemical control on the sugarcane row only; CcWp = chemical control on the whole plot.

Among the six treatments, the weed management was similar on the sugarcane row (i.e. the 75cm-wide soil surface centered on the sugarcane stools) and differed on the inter-row (i.e. the remaining 75cm-wide soil surface between two rows). In all treatments, the sugarcane row was regularly weeded, both chemically and manually. On the inter-row, four treatments (CeE, CeL, DiE, DiL) combined two intercrops, *Desmodium intortum* or *Canavalia ensiformis*, sown either early (between 0 and 2.1 months after sugarcane emergence) or late (between 1.3 and 3.7 months after sugarcane emergence). In a fifth treatment (CcRo), no weeding was carried out on the inter-row. A sixth treatment (CcWp) was a control carried out with chemical weeding of the inter-row; in this control treatment there were no row and inter-row differentiation in weed management (herbicides applications were simultaneous). The CcWp treatment corresponds to the usual practice of sugarcane growers on Reunion.

### 2.3. Intercrop sowing

The date of sowing of the intercrops are summarized in Table 2. Within a trial, the delay between early sowing and late sowing ranged from 1 to 1.6 month. The difference in the date of the early sowing among trials resulted from operational constraints (irrigation availability, mostly). Both *C. ensiformis* and *D. intortum* early sowing treatments in Saint Paul corresponded to a regrowth (natural sowing and regrowth) of the intercrop from a trial carried out the year before (trial which is not included in the study, due to irrigation issues), reason why the sowing date is quoted as ‘0’ in Table 2.

The cover crops were sown manually along two sowing lines 30cm-spaced from each other. On ratooned cane trials, the sugarcane straw present on the soil surface was moved away to allow the sowing of the intercrops and was spread back on the inter row after sowing.

The size of the seeds is very different between both intercrop species, with a mean mass of 1,000 seeds of 1.68 g for *D. intortum* vs. 1,750 g for *C. ensiformis*. The sowing methods were therefore different: *D. intortum* was sown continuously along sowing lines, while *C. ensiformis* was sown in 20cm-spaced poquets containing one seed each. The sowing densities were defined from previous tests: 5.3 kg.ha^-1^ for *D. intortum* and 116.7 kg.ha^-1^ for *C. ensiformis*.

### 2.4. Herbicide applications and manual weeding

When necessary, a pre-emergence herbicide was applied on the cane row and inter-row during the first month after planting or harvesting. The decision to carry out this pre-emergence treatment on the cane rows, as well as the choice of the herbicides, was based on the weed abundance and on the species present in the plot.

In all treatments, post-emergence herbicide applications were carried out on the sugarcane row when the soil coverage by weeds exceeded a 30% threshold on the row. In CeL, DiL and CcWp treatments, post-emergence herbicide applications were carried out on the inter-row before sowing intercrops when the soil coverage by weeds exceeded a 30% threshold on the inter-row. No post emergence herbicide application was carried out on the inter-row after the sowing of the intercrops.

Chemical treatments were carried out with a backpack sprayer with a handheld lance equipped with either a mirror nozzle (average spray rate = 200 L/ha) which allows to spray on a regular step and on a width of 150 or 75 cm, or a slit nozzle (average spray rate = 600 L/ha) which allows to treat by spot on weeds.

Moreover, manual weeding was carried out when needed to discard *Panicum maximum* and *Rottboellia cochinchinensis*, both on rows and inter-rows, in all treatments. Indeed, these two grasses have a high harmfulness on the cane and a strong power of reseeding. As no postemergence herbicides are registered for these weeds, only manual uprooting can control these two species. Lianas were controlled manually in the intercrop treatments and chemically in the control plot when the recovery rate was higher than 30 %.

### 2.5. Plant material

The sugarcane cultivar planted in all trials was ‘R 579’, which is one of the three most common cultivars grown on Reunion. The *C. ensiformis* accession used during our study is registered under the reference CR-XG-00039 in the VATEL germplasm collection managed by Cirad on Reunion (Roux-Cuvelier, 2017). The *C. ensiformis* seed were multiplied on an experimental station field to supply the trials. The *D. intortum* seeds, cultivar ‘Greenleaf’, were imported from Australia (Williams Group, Murwillumbah, Australia) and purchased from a local seed retailer (Association Réunionnaise de Pastoralisme, Plaine des Cafres, Reunion). Germination tests were carried out on the seed batches prior sowing; only seed batches exhibiting a germination rate higher than 90% were used.

### 2.6. Field measurements

To quantify the herbicide application in each treatment in each trial, the Treatment Frequency Index or TFI (Roβberg et al. 2002, cited by Sattler et al. 2007; Brunet et al. 2008) was computed. The TFI is an indicator used to assess the use of pesticides on different cropping systems. TFI has an additive construction, that is, TFI increases with the number pesticide applications and the dosage, but does not consider the compound toxicity (Simon et al. 2011). For each treatment, the TFI was calculated as follows:

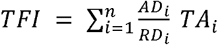

where n is the number of herbicide applications, AD_j_ the actual dosage sprayed of the compound, RD_i_ the lowest recommended dosage of the compound, and TA_i_ the proportion of the plot area sprayed (0< TA <1). To compute the TFI, the sugarcane row was accounted as TA=0.5 and the inter-row as TA=0.5, as exposed above. Details about the herbicide compounds used and their recommended dosages are in Supplementary Table S2. The actual dosage *AD*_*j*_ was computed proportionally after measuring the remaining of the mixture in the sprayer tank after spraying.

Cane yield was measured at harvest. The stalks from the two central rows were manually harvested and weighted (i.e. an harvested area of 30 m^2^ per plot).

Ground cover by the weeds (considered as a whole) or by the intercrops was assessed based on a visual notation (Marnotte et al. 2001). The visual notation was carried out on the interrow and on the whole plot separately. The weed species present on each plot were recorded, except in the Saint Benoît 2013-2014 trial. Depending on the trial and the crop season, 7 to 8 visual notations were carried out during 10 to 14 months after sugarcane emergence.

Stalk borer damage assessment was performed just before harvest. 30 stalks (three segments of 10 consecutive stalks) were harvested in the two central rows of each plot and dissected for damage assessment. A stalk was considered damaged when at least one internode was bored by *C. sacchariphagus* larvae.

### 2.7. Statistical analysis

Due to the failure of the *D. intortum* intercrop installation in some trials (see the Results section), the two treatments with *D. intortum* were discarded from the dataset prior to the statistical analysis.

Because the TFI data and the number of manual weeding were computed for each treatment at the trial level and not at the elementary plot level, these two variables were analyzed with a general linear model, considering the trial x crop cycle as a fixed block effect (i.e with no intra trial replications).

The ground coverage and yield data were analyzed using a linear mixed model. The treatment, replication, location, treatment x location were considered as fixed effects. The correlation of model residuals due to the longitudinal observations (i.e. two trials were observed during two successive crop cycles) was taken into account with a ‘repeated’ instruction in SAS procedure MIXED (SAS Institute Inc. 2023). We compared different covariance structures (variance components or compound symmetry) and chose the most appropriate one based on the model fit estimated by the AIC.

The proportion of borer damaged stalks was analyzed in a similar way but using a generalized linear mixed model with SAS procedure GLIMMIX with a binomial distribution of the data and a logit link (SAS Institute Inc. 2023).

## 3. Results

### 3.1. Intercrop installation and ground cover

Whether sown on early or late, *C. ensiformis* quickly covered the inter-row (Figure 1). There was a quasi-disappearance of *C. ensiformis* at sugarcane harvest time. On the other hand, ground cover by *D. intortum* was lower than *C. ensiformis* (Figure 1). Ground cover was very low or nil on half of the trials x crop cycle, justifying the discarding of the *D. intortum* treatments from the rest of the analysis presented hereafter.

**Figure 1.**
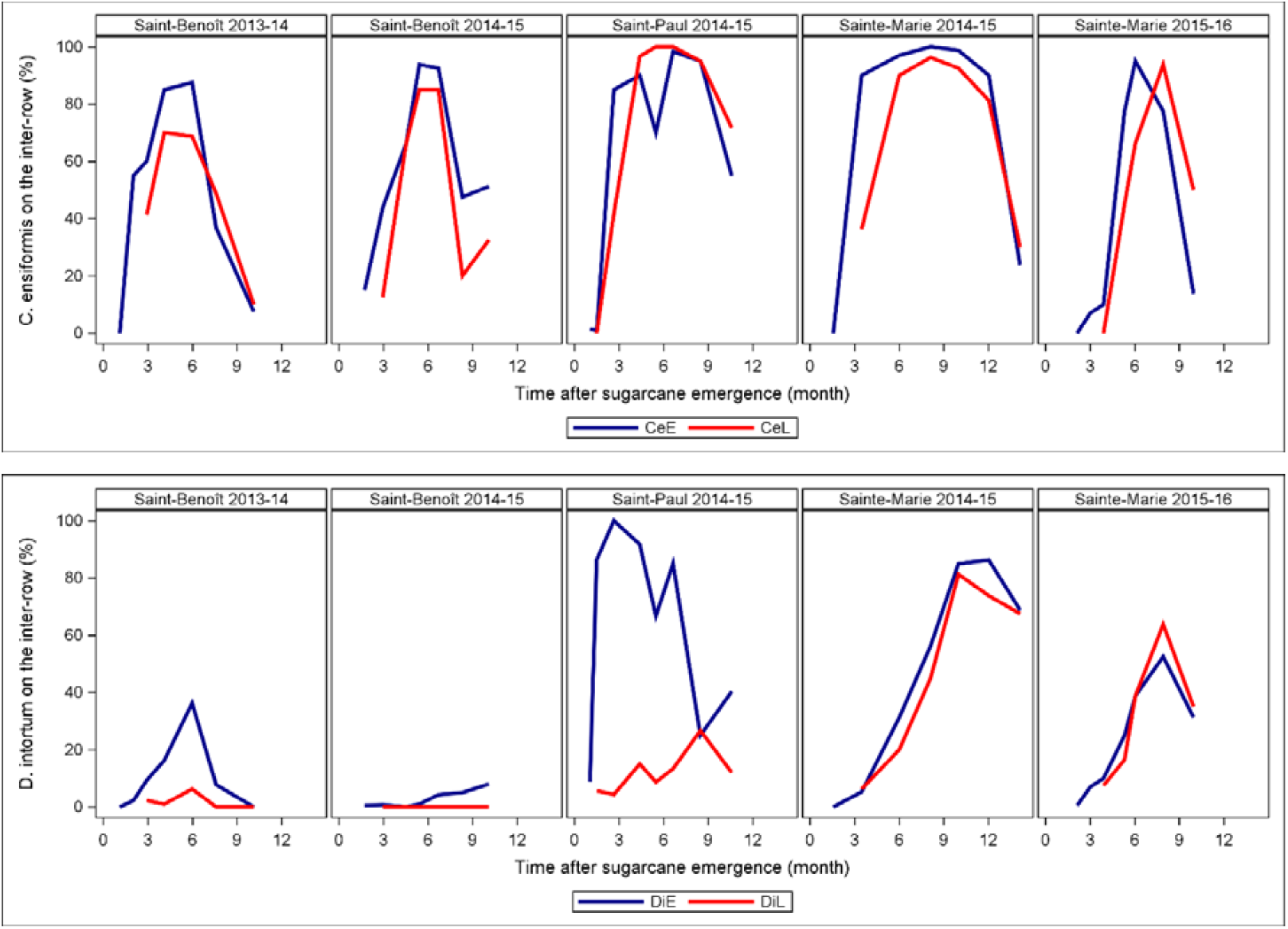
Evolution of the mean ground coverage by C. ensiformis orD. intortum intercrops in the inter-row. CeE = C. ensiformis sown early; CeL = C. ensiformis sown late. DIE= D. intortum sown early; DIL = D. intortum sown late.

In the untreated inter-rows (CcRo treatment), 53 weed species were observed across the three locations (Supplementary Table S3). The most abundant weed species in each trial and crop season are listed in Supplementary Table S4.

### 3.2. Manual and chemical weeding

The mean treatment frequency index TFI was significantly different between treatments (Table 3). The two *C. ensiformis* intercrop treatments significantly reduced the TFI by 28% and 63%, for CeL and CeE respectively, compared to the chemical control on the whole plot (CcWp). The TFI of the CeE treatment was significantly lower than CeL. The TFI of both *C. ensiformis* intercrop treatments did not differ significantly from the chemical control on the row CcRo treatment.

**Table 3.**
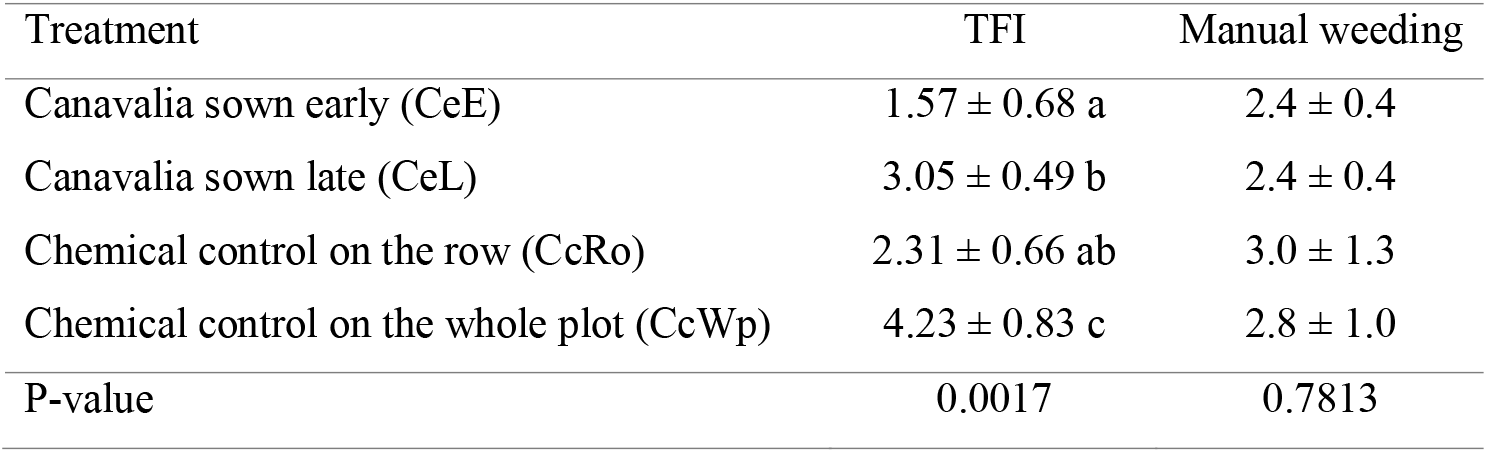
Comparison of the treatment frequency index (TFI) and the number of manual weeding between the treatments. Means (± SEM) followed by different letters are significantly different (P < 0.05) according to pairwise tests of least square mean differences with Tukey adjustment for multiple comparisons.

The mean number of manual interventions was not significantly different between treatments.

### 3.3. Ground coverage by weeds, cane yield and stalk borer damage

The mean ground coverage on the inter-row by both Canavalia and weeds was significantly higher on the CeE and CeL treatments than on the naturally weeded inter-row treatment (CcRo) (Table 4). The mean ground coverage was increased by a 1.8-fold factor on the early sown Canavalia (CeE), and by a 1.6-fold factor on the late sown Canavalia (CeL). Both Canavalia intercrop treatments allowed a reduction of soil mean coverage by weeds compared to the chemical control on the whole plot (CcWp), only the reduction by CeL being significant.

**Table 4.** Comparison of the cane yield and stalk borer damages between the treatments. Means (± SEM) followed by different letters are significantly different (P < 0.05) according to pairwise tests of least square mean differences with Tukey adjustment for multiple comparisons.

| Treatment | Mean ground coverage by weeds and Canavalia on the inter-row (%) | Mean ground coverage by weeds on the inter-row (%) | Yield (t.ha<sup>-1</sup>) | Bored stalks (%) |
| --- | --- | --- | --- | --- |
| Canavalia sown early (CeE) | 62.8 ± 3.3 c | 7.3 ± 1.5 ab | 128.6 ± 5.1 b | 52.0 ± 2.4 a |
| Canavalia sown late (CeL) | 54.2 ± 2.9 c | 7.5 ± 1.9 a | 140.5 ± 4.2 ab | 58.9 ± 2.3 ab |
| Chemical control on the row (CcRo) | 34.8 ± 3.9 b | 34.8 ± 3.9 c | 139.2 ± 5.9 ab | 54.0 ± 2.3 ab |
| Chemical control on the whole plot (CcWp) | 17.1 ± 2.8 a | 17.1 ± 2.9 b | 144.0 ± 3.8 a | 68.4 ± 2.2 b |
| p-values |  |  |  |  |
| Treatment | < 0.0001 | < 0.0001 | 0.0433 | 0.0376 |
| Locality | 0.0386 | 0.1819 | < 0.0001 | 0.1144 |
| Treatment x Locality | 0.5046 | 0.0988 | 0.3500 | 0.4073 |

Sugarcane yields ranged from 129 to 144 t.ha^-1^ and were significantly influenced by treatment and location (Table 4). The CeE (Canavalia sown early) treatment exhibited a significant yield loss estimated by the model at 18.6 ± 6.3 t.ha^-1^ when compared with the control treatment CcWp (chemical control on the whole plot). The two other treatments, CeL (Canavalia sown late) and CcRo (unweeded inter-rows) did not differ significantly from the CcWp control treatment.

The stalk borer damage was significantly influenced by treatment but not by locality (Table 4). A significant reduction of borer damage by a 0.8-fold factor occurred in the Canavalia early sown treatment (CeE) compared to the chemical control (CcWp). The three other treatments did not differ significantly from each other.

## 4. Discussion

During this study, we have demonstrated that the use of *C. ensiformis* intercrop, installed between 0 to 3.7 month after sugarcane emergence and combined with manual weeding and a reduced number of herbicide sprays, allowed a control of weed populations equivalent or better than a pure chemical control of weeds. The reduction of herbicide use allowed a decrease of the TFI ranging from 28% to 63%, without causing an increase of the number of manual weeding interventions. However, we also observed in our trials that an early *C. ensiformis* intercrop installation (by natural sowing or regrowing) caused a significant yield loss of 18.6 t.ha^-1^. Additionally, we observed a reduction of the percentage of stalks damaged by the borer *C. sacchariphagus* associated with an early installation of the *C. ensiformis* intercrop. Finally, we observed that *D. intortum* failed to develop in half of the situations studied.

In conclusion, this study illustrates the promising use of intercropping the sugarcane with *C. ensiformis* to reduce weed populations and to allow a reduction of herbicide use. However, it also appeared in this study that an early installation of the intercrop can result in competition with the sugarcane crop and cause a yield loss. On the other hand, we observed that the late sowing of *C. ensiformis*, in possible combination of a pre-emergence herbicide treatment, remains an interesting solution, showing no adverse impact on sugarcane yield in our trials and allowing a reduction of herbicide use of 28%.

It is unclear how the reduction in stem borer damage was associated solely with the early intercropping of *C. ensiformis*. Several hypotheses are indeed often invoked to explain the beneficial effect of intercropping on pest control, among which a repellant effect of the intercrop (Khan et al. 2001), a diversion effect which disrupts the visual stimuli used by the pest to identify its host plant (Finch & Collier, 2000), or an increase of the populations of the natural enemies of the pest by providing alternative food sources like nectar of alternative preys (Nikpay et al. 2023). Here, none of these hypotheses appear satisfactory, as none can explain how no damage reduction was observed in the late sowing *C. ensiformis* intercrop. One remaining hypothesis is that the significant reduction of sugarcane biomass caused by the intercrop could have resulted in a reduction of attractivity of the crop for the stalk borer ovipositing females.

## Acknowledgements

This work was co-funded by the French Ministry of Agriculture (CASDAR grant 1269), the European Union’s Agricultural Fund for Rural Development, the Conseil Départemental de La Réunion, the Conseil Régional de La Réunion and the Centre de Coopération Internationale en Recherche Agronomique pour le Développement. We would like to thank Pascal Marnotte for his useful advices, and also Richard Tibère, and Cédric Lallemand for their technical help. We would also like to thank the sugarcane growers involved in the field experiments.

## Supplementary material

Raw data are available at https://doi.org/10.18167/DVN1/NVW8ZV

SAS codes used for the statistical analysis are available at https://gitlab.cirad.fr/pvbmt/2025_biorxiv_canavalia_intercrop_sugarcane

