## Supplementary material for "*Canavalia ensiformis* intercrop to reduce weeds and stalk borer damage in sugarcane"

*Supp. Table S1. Fertilization and harvest date of the trials.*

| Trial | Fertilization | Harvest date |
| --- | --- | --- |
| Sainte Marie 2014-2015 | Two split applications :   - 410 kg/ha 18-7-30 - 200 kg/ha 18-7-30 | 3-5th August 2015 |
| Sainte Marie 2015-2016 | Two split applications :   - 600 kg/ha 15-12-24 - 300 kg/ha 15-12-24 + 70 kg/ha Urea 46% | 1-4th August 2016 |
| Saint Paul 2014-2015 | Two split applications :   - 650 kg/ha 20-9-27 - 285 kg/ha 20-9-27 + 217 kg/ha Urea 46 % | 16-19th October 2015 |
| Saint Benoît 2013-2014 | Two split applications :   - 600 kg/ha 9-15-22 - 200 kg/ha 18-7-30 | 8-15 july 2014 |
| Saint Benoît 2014-2015 | Two split applications :   - 600 kg/ha 18-7-30 - 300 kg/ha 18-7-30 | 9-13 july 2015 |

*Supp. Table S2. Characteristics of the herbicide compounds used in the trials.*

| Compound composition (active ingredients and concentration) | Compound’s lowest Recommended Dose  (RD) | Period of application | Weed targets |
| --- | --- | --- | --- |
| S-metolachlor 400g/L  mesotrione 40 g/L  benoxacor 20 g/L | 3.75 L/ha | pre-emergence and early post-emergence | small poacea and some dicotyledons |
| isoxaflutole 750 g/kg | 0.133 kg/ha | pre-emergence | some poacea (Panicum maximum) |
| pendimethalin 400 g/L | 3 L/ha | pre-emergence | some poacea (*Rottboellia cochinchinensis*) |
| metribuzin 700 g/kg | 1,25 kg/ha | pre-emergence and post-emergence | small poacea and some dicotyledons |
| mesotrione 100 g/L | 1,5 L/ha | post-emergence | dicotyledons (in addition to other herbicides) |
| 2,4-D 600 g/L | 2 L/ha | post-emergence | some dicotyledons at a young age |
| fluroxypyrl ester 1-methylheptyl 200 g/L | 1 L/ha | post-emergence | lianas and other dicotyledons |
| glufosinate ammonium 150 g/L | 5 L/ha | post-emergence (general treatments) | most weeds |
| glyphosate 480 g/L | 5 L/ha | post-emergence (general treatments) | most weeds |

*Supp. Table S3 List of weed species observed on the inter row of the CcRo treatment (CcRo = chemical control on the sugarcane row only) in four trials (no data collected in the Saint Benoît 2013-2014 trial).*

| Species | Species (continued) |
| --- | --- |
| *Ageratum conyzoides Amaranthus dubius Argemone mexicana Asystasia gangetica Bidens pilosa Cajanus scarabaeoides Cardiospermum halicacabum Centrosema pubescens Chamaesyce hirta Clidemia hirta Commelina benghalensis Commelina diffusa Conyza sumatrensis Crassocephalum crepidioides Crotalaria retusa Croton bonplandianus Cyclospermum leptophyllum Cynodon dactylon Cyperus rotundus Cyperus* sp. *Desmanthus virgatus Digitaria horizontalis Digitaria* sp. *Emilia sonchifolia Euphorbia heterophylla Indigofera hirsuta* | *Ipomoea hederifolia Ipomoea obscura Lantana camara Litsea glutinosa Lycopersicon esculentum  Panicum maximum (Megathyrsus maximus) Mimosa pudica Mirabilis jalapa Momordica charantia Oxalis corniculata Paederia foetida Paspalum scrobiculatum Paspalum sp. Passiflora foetida Phyllanthus niruroides Rottboellia cochinchinensis Senna occidentalis Setaria barbata Sida acuta Sida glabra Sida retusa Sigesbeckia orientalis Solanum lycopersicum Solanum mauritianum Solanum nigrum Tephrosia purpurea Teramnus labialis* |

*Supp. Table S4. List of the weed species contributing together to at least 90% of the mean soil coverage of the CcRo treatment (CcRo = chemical control on the sugarcane row only) in each trial. Species are sorted by decreasing order of mean soil coverage within each trial. No data collected in the Saint Benoît 2013-2014 trial.*

| Saint Paul 2014-2015 | Saint Benoît 2014-2015 | Sainte Marie 2014-2015 | Sainte Marie 2015-2016 |
| --- | --- | --- | --- |
| *Setaria barbata Cyperus rotundus Ageratum conyzoides Oxalis corniculata Solanum nigrum Rottboellia cochinchinensis* | *Cynodon dactylon Clidemia hirta Ageratum conyzoides Litsea glutinosa Rottboellia cochinchinensis Cyperus rotundus Solanum nigrum Commelina diffusa Oxalis corniculata* | *Croton bonplandianus Centrosema pubescens Ipomoea obscura Solanum nigrum Mimosa pudica Argemone mexicana Oxalis corniculata Senna occidentalis Indigofera hirsuta Amaranthus dubius Crotalaria retusa Bidens pilosa Ageratum conyzoides Chamaesyce hirta Commelina benghalensis Cyperus rotundus Solanum lycopersicum* | *Mimosa pudica Ipomoea obscura Cyperus rotundus Centrosema pubescens Croton bonplandianus Cardiospermum halicacabum Senna occidentalis Cajanus scarabaeoides Argemone mexicana Bidens pilosa Indigofera hirsuta Tephrosia purpurea* |

*Supp. Figure S1 Evolution of the mean soil coverage by weeds and by C. ensiformis (%) on the whole plot or on the inter-row in the five field trials. Green = chemical control on the sugarcane row only (CcRo), red = C. ensiformis sown late (CeL), orange = chemical control on the whole plot (CcWp), blue = C. ensiformis sown early (CeE).*

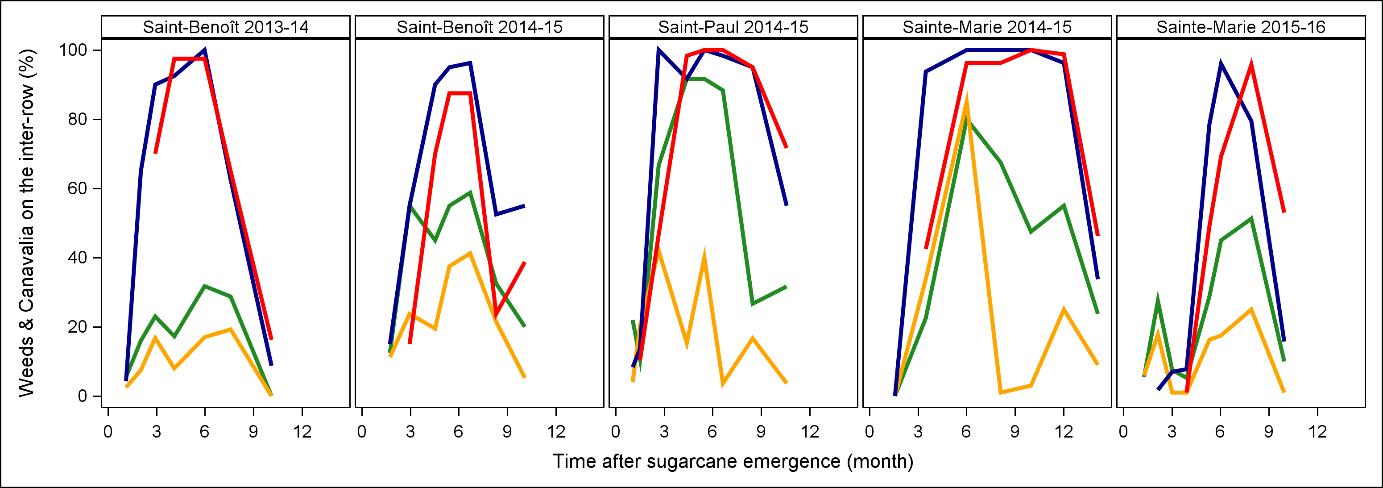

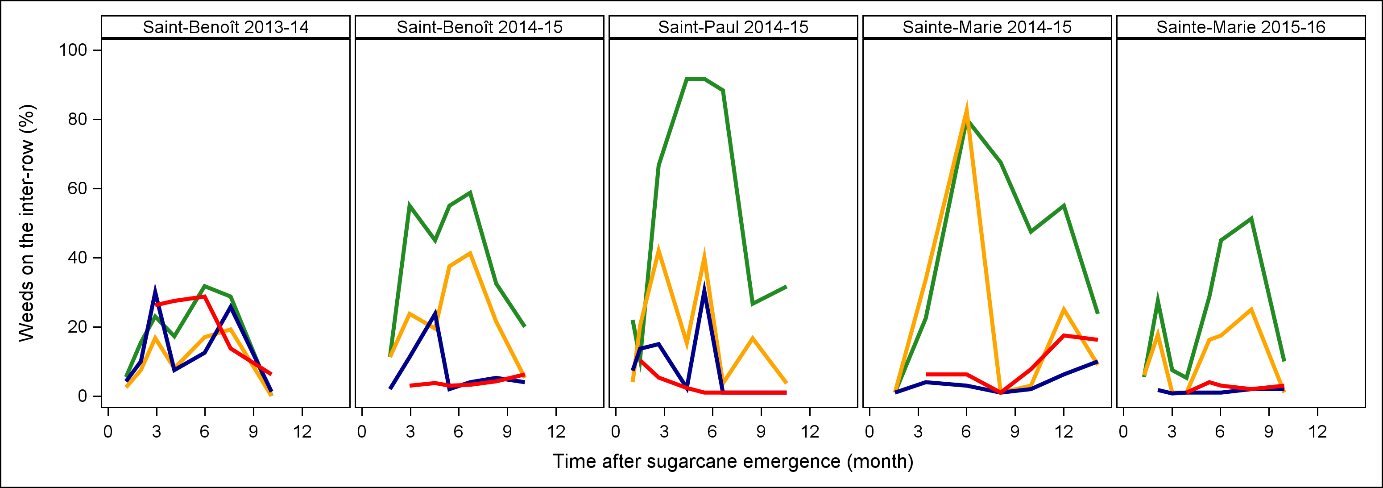

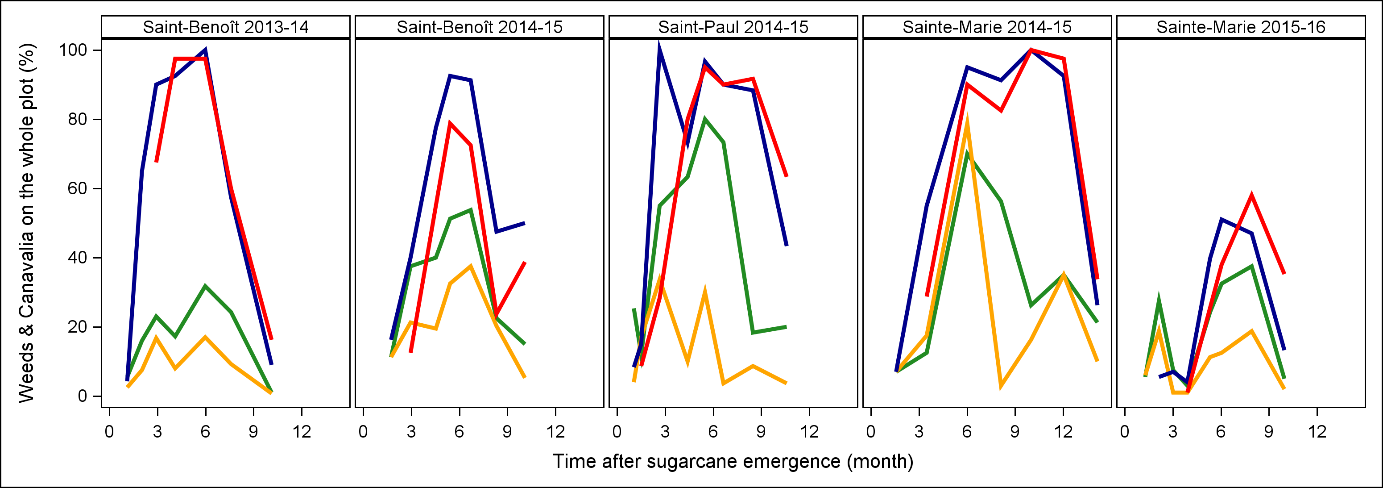

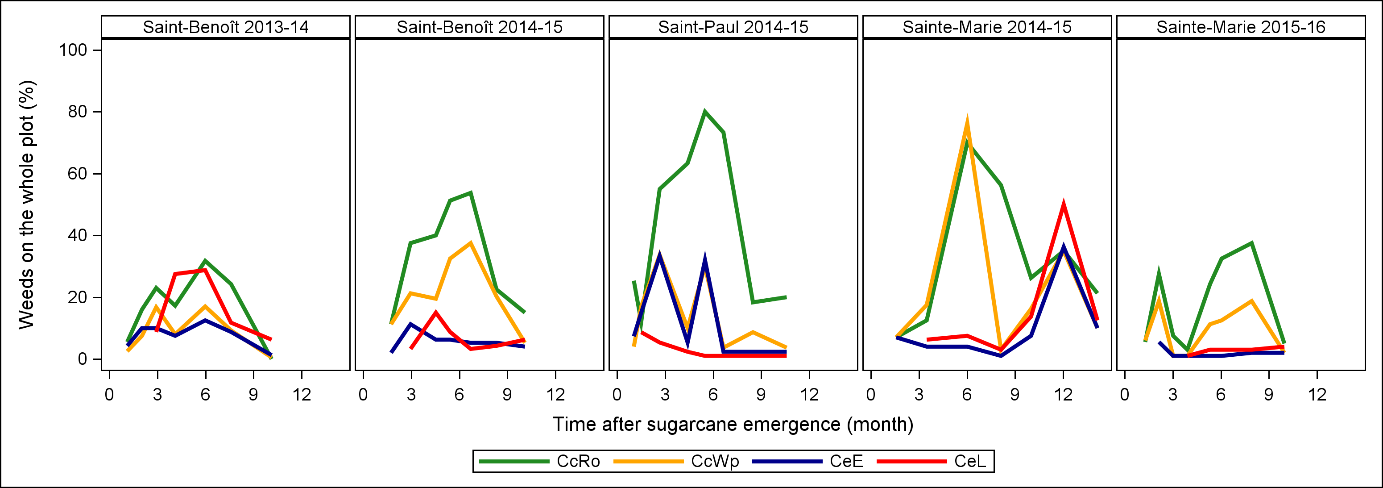
